# Thyroid hormone regulates proximodistal identity in the fin skeleton

**DOI:** 10.1101/2020.08.18.256354

**Authors:** Yinan Hu, Melody Harper, Benjamin Acosta, Joan Donahue, Hoa Bui, Hyungwoo Lee, Stacy Nguyen, Sarah McMenamin

## Abstract

Across the ∼30,000 species of ray-finned fish, fins show incredible diversity in overall shape and in the patterning of the supportive bony rays. Fin length mutant zebrafish have provided critical insights into the developmental pathways that regulate relative fin size. However, the processes that govern skeletal patterning along the proximodistal axis of the fin have remained less well understood. Here, we show that thyroid hormone regulates proximodistal identity of fin rays, distalizing gene expression profiles, morphogenetic processes during outgrowth, and ultimate morphology of the fin. This role for thyroid hormone in specifying proximodistal identity appears conserved between development and regeneration, in all the fins, and between species. We demonstrate that proximodistal identity is regulated independently from pathways that determine size, and we show that modulating proximodistal patterning relative to growth can recapitulate the spectrum of fin ray diversity found in nature.

## Introduction

How organisms create form is a fundamental question in biology (Thompson, 1942; Gould, 1977). The processes that determine size and pattern must be closely coordinated during development, and relative changes in these processes can profoundly change form (e.g. by altering number of body segments; Gomez and Pourquié, 2009). Organ growth and tissue morphogenesis can be cued by local signaling factors, voltage gradients and cell-to-cell interactions (reviewed in Heller and Fuchs, 2015; Tung and Levin, 2020). In addition to these local processes, global factors such as hormones can coordinate morphogenesis in disparate tissues, and such systemic signals may be leveraged to establish or reinforce patterns within a complex organ.

Fins are phenomenally diverse across teleosts. These propulsive structures share deep homologies with tetrapod limbs (Schneider and Shubin, 2013; Nakamura et al., 2016), and the fin-to-limb transition involved major changes in patterning and proportion along the proximodistal axis (e.g. see Woltering et al., 2020). Fin rays show distinct proximodistal polarity: ray segments taper and shorten distally, periodically bifurcating into branches (Christou et al., 2018). This patterning effectively determines the biomechanical properties of the fin (Aiello et al., 2018; Puri et al., 2018), and varies considerably across species, from uniformly thick rays with no branches (e.g. sygnathids, sculpins), to tapering rays with numerous branches (e.g. guppies, killifish).

Bioelectric signaling regulates the size of fins (Iovine et al., 2005; Perathoner et al., 2014; Stewart et al., 2019; Harris et al., 2020), but the mechanisms underlying proximodistal patterning have remained largely elusive. Sonic hedgehog (Shh) is essential in branch formation (Laforest et al., 1998; Zhang et al., 2012), and the local morphogenesis establishing bifurcations is now emerging (Armstrong et al., 2017; Braunstein et al., 2020). However, it remains unknown what proximodistal identity cues allow branching machinery to be deployed at the correct locations along growing rays. Retinoic acid (RA) is a classic mediator of proximal identity in tetrapods (Maden, 1982; Tickle et al., 1982; Yashiro et al., 2004), and exogenous RA appears to change the relative location of fin ray bifurcations (White et al., 1994). However, this is likely due to fusion of sister rays rather than a genuine change in proximodistal identity (White et al., 1994; Blum and Begemann, 2012). Furthermore, while some pathways governing segment length have been discovered (Sims Jr et al., 2009; Schulte et al., 2011), mechanisms governing the progressive distal shortening of segments remain unknown (although they can be modelled computationally; Rolland-Lagan et al., 2012).

Thyroid hormone (TH) is an essential regulator of cellular metabolism and homeostasis, and coordinates developmental events in diverse vertebrates (McMenamin and Parichy, 2013; Shi, 2013; Buchholz, 2015). In zebrafish, the hormone regulates numerous aspects of post-embryonic development (McMenamin et al., 2014; McMenamin et al., 2017; Hu et al., 2019; Keer et al., 2019; Saunders et al., 2019). In hypothyroid zebrafish, we observed changes in the fin skeleton consistent with shifts in axis polarity, and we hypothesized that TH regulates proximodistal identity of the fin rays.

## Results and Discussion

### TH promotes distal identity in *Danio* fin rays

Changes to developmental TH titer dramatically changed the proximodistal position of bifurcations in adult fins (Figure 1A-D). More subtle aspects of proximodistal morphology of the rays also showed dependence on TH: distal segments from hypothyroid backgrounds remained longer and retained higher bone density than those in euthyroid backgrounds (Figure 1E-F; Figure 1—Figure Supplements 1-2). This role for TH in promoting distal features appeared consistent in all paired and medial fins (Figure 1—Figure Supplement 3), in a congenitally hypothyroid zebrafish mutant, and in another species, *D. albolineatus* (Figure 1—Figure Supplement 4).

**Figure 1.**
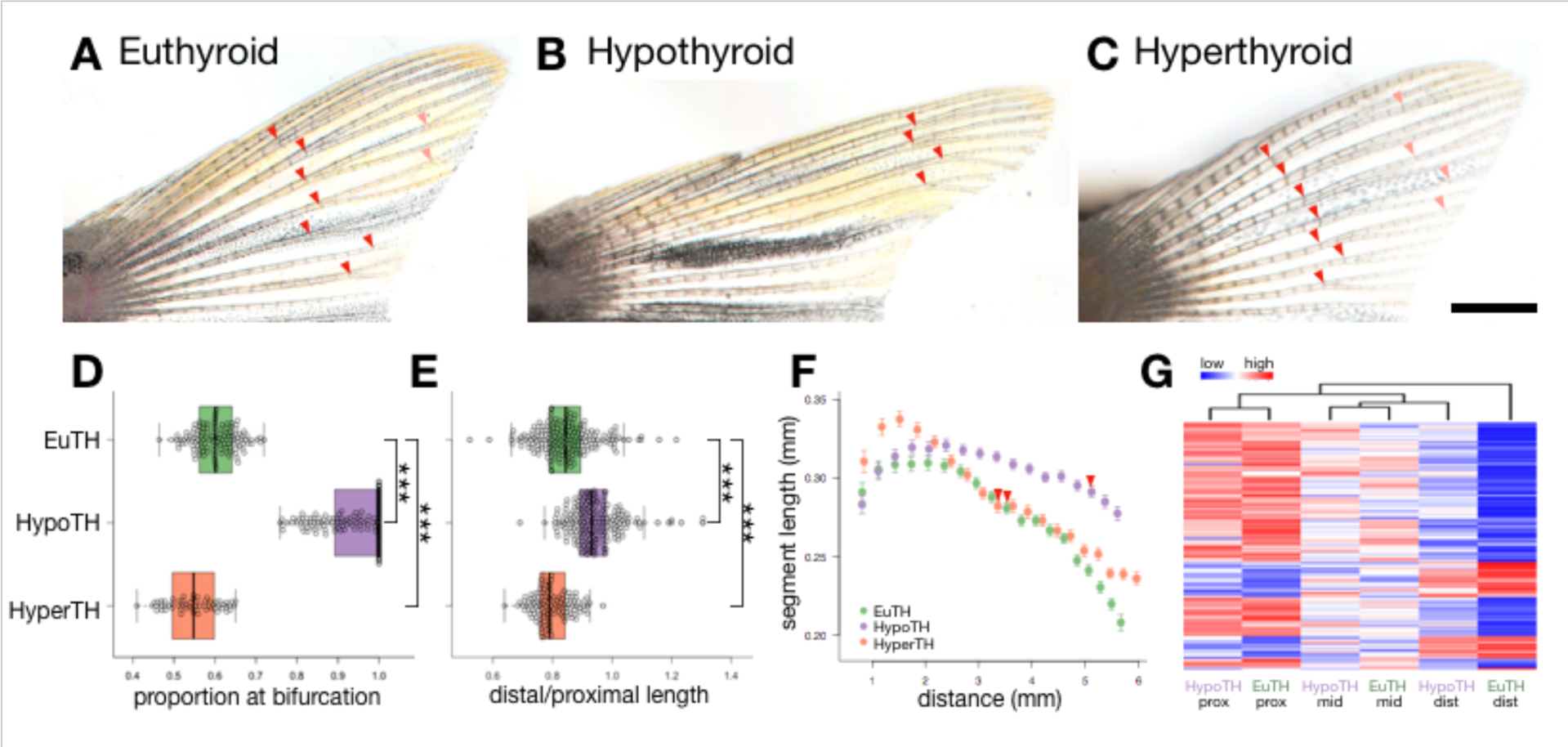
TH promotes distal identity in fin rays. (**A-C**) Dorsal lobes of caudal fins from zebrafish reared under different TH profiles. Red and pink arrowheads indicate primary and secondary bifurcations, respectively. Scale bar, 1 mm. (**D**) Proportion of the total ray length at the primary bifurcation. (**E**) Ratio of distal segment length (15-17^th^ segments) to proximal segment length (5-7^th^ segments). Significance determined by ANOVA and Tukey’s HSD. (**F**) Plot showing the average length of each segment (starting from 3^rd^ segment) and distance from the segment to the body. Whiskers represent 95% confidence intervals. Arrowheads indicate the average location of the primary bifurcation in each background. (**G)** Heatmap of 128 genes expressed in a proximodistal gradient that shows dependenence on TH. Dendrogram at top represents hierarchical clustering of the samples; note that distal regions from hypothyroid cluster with middle regions from euthyroid. EuTH, euthyroid; HypoTH, hypothyroid; HyperTH, hyperthyroid.

Many genes are expressed in a gradient along the proximodistal axis of the adult caudal fin (Rabinowitz et al., 2017), and we found that the majority of such gradients (128/233; 55%) were dependent on the presence of TH (Figure 1G; Figure 1—Figure Supplement 5). In the absence of TH, fins showed proximalized expression profiles, particularly in distal regions. With one exception (*fabp1b*.*2*), RA-pathway genes maintained similar proximodistal gradients and expression patterns under hypothyroidism (Figure 1—Figure Supplement 5), suggesting TH does not mediate proximodistal identity by regulating expression of the RA pathway.

### Proximodistal identity is independent from the pathways that regulate size

Although TH regulates distal morphology and expression, the hormone surprisingly had little influence on fin area or ray length (Figure 1—Figure Supplement 1G-I). Further, hypothyroidism proximalized fins of *longfin* and *shortfin* mutants (Figure 2A-F; Figure 2—Figure Supplement 1), suggesting that TH functions independently from the bioelectricity pathways altered in these fin length mutants. More broadly, we conclude that the mechanisms regulating proximodistal patterning are distinct from those that regulate overall size, and posit that these processes may be decoupled during development and evolution.

**Figure 2.**
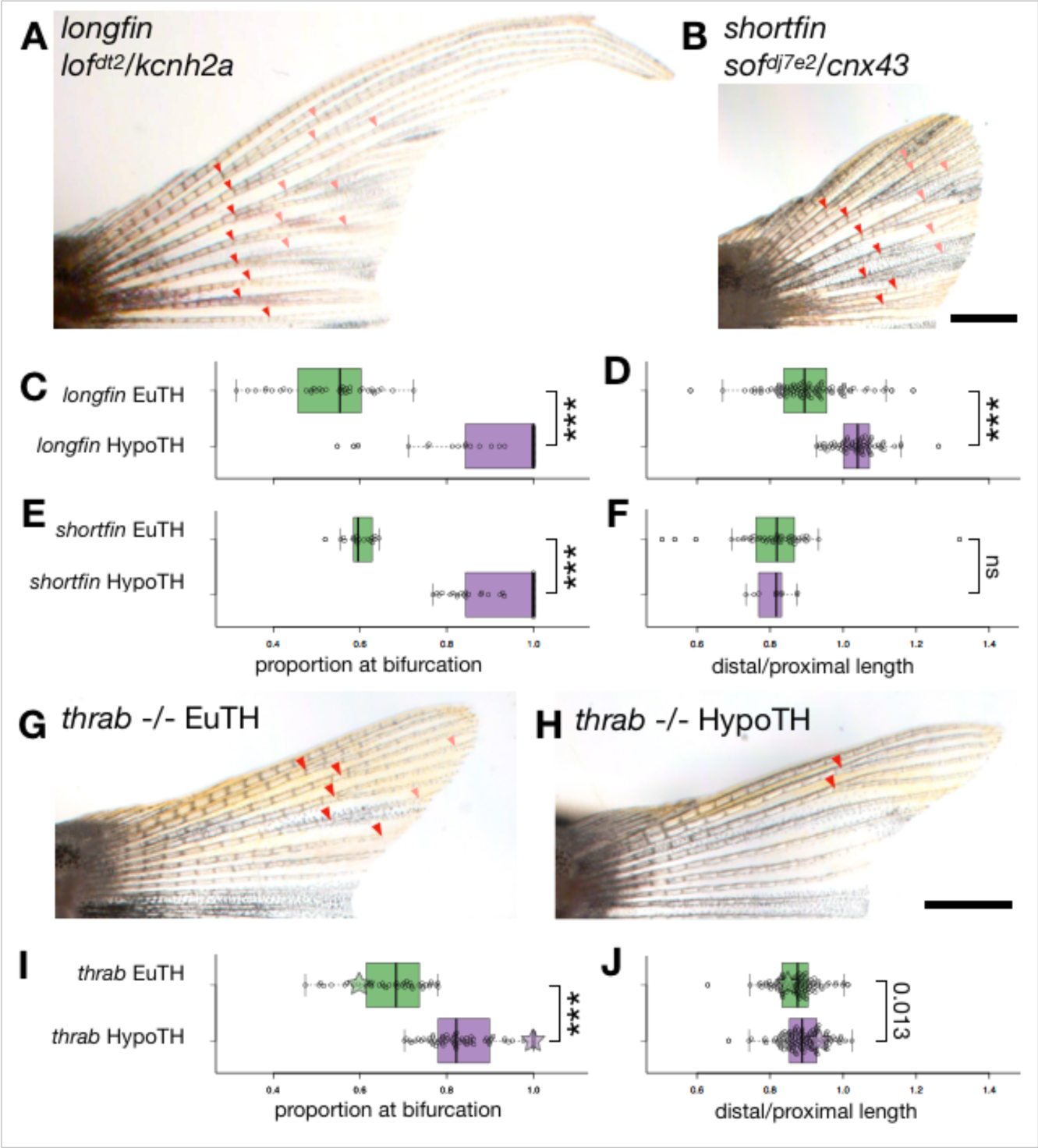
Bioelectricity pathways determine fin length independent of TH, and TH receptor positively and negatively regulates distal identity. Caudal fins of (**A**-**B)** *longfin* and *shortfin* mutants and (**G-H**) *thrab* -/-mutants reared under (G) euthyroid or (H) hypothyroid conditions. Arrowheads, bifurcations; Scale bars, 1 mm. Box plots showing (**C-E** and **I**) proportion of the total ray length at the primary bifurcation and (**D-F** and **J**) ratio of distal segment length to proximal segment length. Significance determined by ANOVA and Tukey’s HSD. Green and purple stars overlaid in I and J indicate the mean values for non-mutant reared under eu- and hypothyroid conditions.

To explore the phenotypic diversity that might result from changes to proximodistal identity, we built a simple computational model to simulate fin ray morphology based on two input parameters. By adjusting “distalization,” we produced rays resembling those of TH-disrupted fish; adjusting “size,” we generated rays resembling those of fin length mutants (Figure 2—Figure Supplement 2). Further modulating distalization to its extremes, we could simulate rays resembling those of other species (Figure 2—Figure Supplement 2), potentially capturing modular developmental changes that may underlie adaptation and evolved diversity.

### Thrab positively and negatively shapes proximodistal identity

TH primarily acts through dual-action nuclear TH receptors (Hörlein et al., 1995). We found that euthyroid *thrab* mutants showed moderately proximalized fin phenotypes relative to euthyroid wild-type, suggesting that liganded Thrab promotes distalization (Fig. 2G). Unliganded TH receptors serve to repress developmental progress in other contexts (Buchholz et al., 2003; Saunders et al., 2019), and we asked if Thrab contributed to proximodistal identity in the absence of TH. Indeed, rays of hypothyroid *thrab* mutants showed partially rescued distal morphologies relative to non-mutant hypothyroid fish (Figure 2 G-J; Figure 2—Figure Supplement 1). We conclude that liganded Thrab promotes distalization, while unliganded Thrab represses distalization.

### TH distalizes rays during outgrowth

Treatment with TH throughout regeneration is sufficient to robustly rescue distal identity (Figure 3B). Previous studies have shown that cellular memory within the blastema dictates fin size during regeneration (Tornini et al., 2016; Wang et al., 2019), and we asked if the TH rescue of distal morphology was due to effects of the hormone on the blastema. We treated hypothyroid fish with TH (in the form of T4) for 24 hours during each of the first 3 days post amputation (dpa), but found that none of these treatments were sufficient to rescue distal patterning in regenerates (Figure 3—Figure Supplement 1). This led us to hypothesize that TH may be acutely required to specify distal identity, and we tested this possibility by delaying TH treatments until late stages of regenerative outgrowth. Euthyroid fish generally form clearly separated bifurcations by 7 dpa, but even a T4 rescue starting at 14 dpa was able to rescue branching and distal morphology (Figure 3C; Figure 3—Figure Supplement 2C-F). Instead of having a critical window for TH sensitivity, the fin rays remain competent to become distalized by TH throughout regeneration.

**Figure 3.**
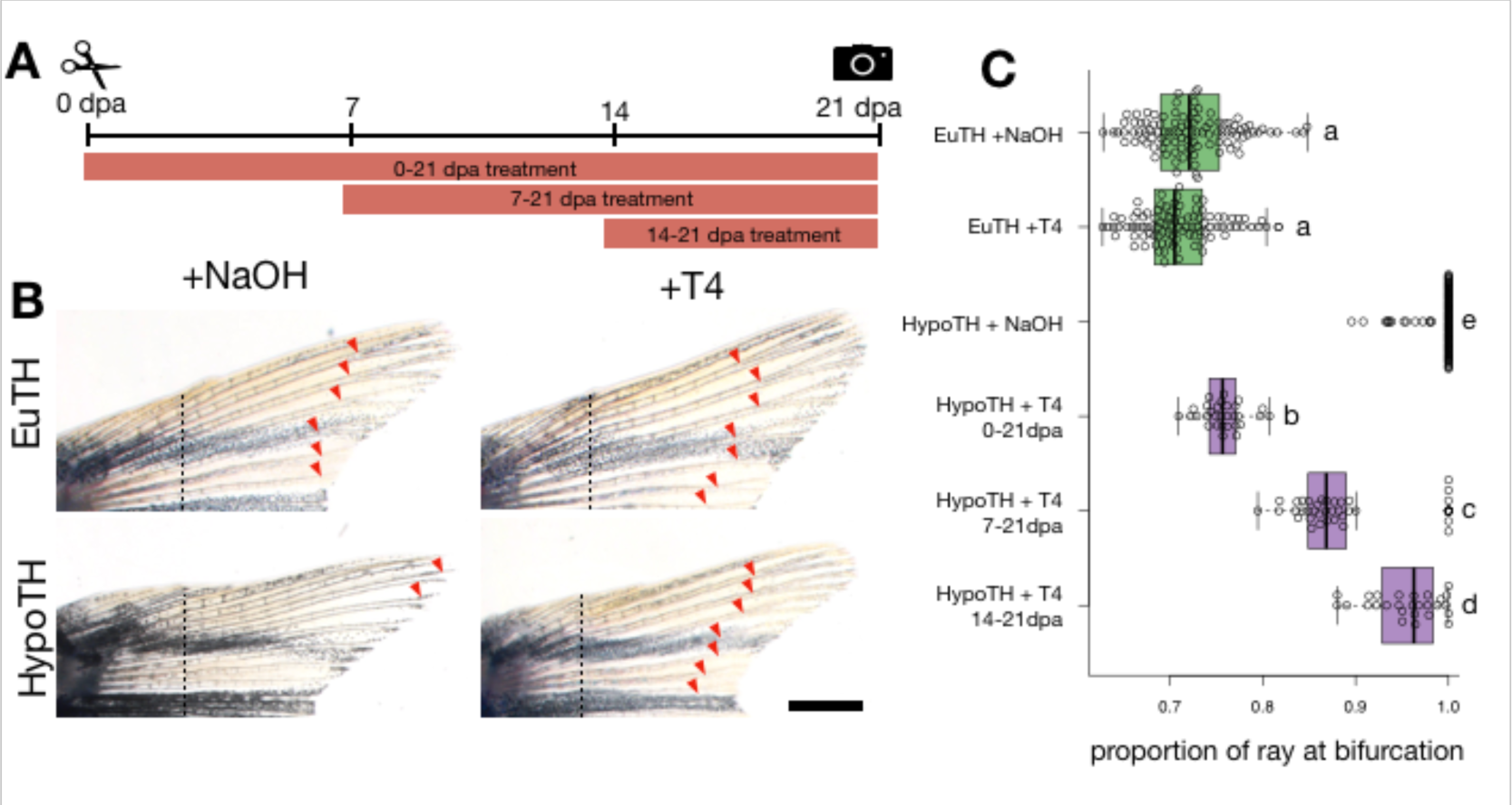
Exogenous TH is sufficient to distalize hypothyroid fin rays during regeneration. (**A**) Experimental timeline. (**B**) Fins from euthyroid or hypothyroid fish treated with NaOH vehicle or 10 uM T4 throughout regeneration (0-21 dpa). Dashed lines indicate planes of amputation. Arrowheads, bifurcations; Bar, 1 mm. (**C**) Proportion of total ray length at bifurcation for fins of eu- and hypothyroid fish regenerated under different treatment profiles. Timing of treatment did not alter proportion for the euthyroid groups or for the NaOH-treated hypothyroid control, and these groups are shown here pooled; also see Figure 3—Figure Supplement 2D.

### TH induces branching morphogenesis

We found that small domains of TH activity were present at the distal tips of both uninjured and regenerating rays (Figure 4A; Figure 4—Figure Supplement 1). Tissue adjacent to the lateral-most rays also showed high TH activity, and removal of the lateral rays did not cause the next-most lateral rays to adopt high TH activity (Figure 4— Figure Supplement 1B-E). Tips of growing rays showed an increase in TH activity preceding branch formation (Figure 4—Figure Supplement 1F), potentially reflecting local regulatory signaling or elevated metabolic demands during bifurcation. Tips of regenerating rays showed *thrab* expression (Figure 4B), but different proximodistal levels of fin regeneration showed no differences in the amount of *thrab* expression (Figure 4—Figure Supplement 2).

**Figure 4.**
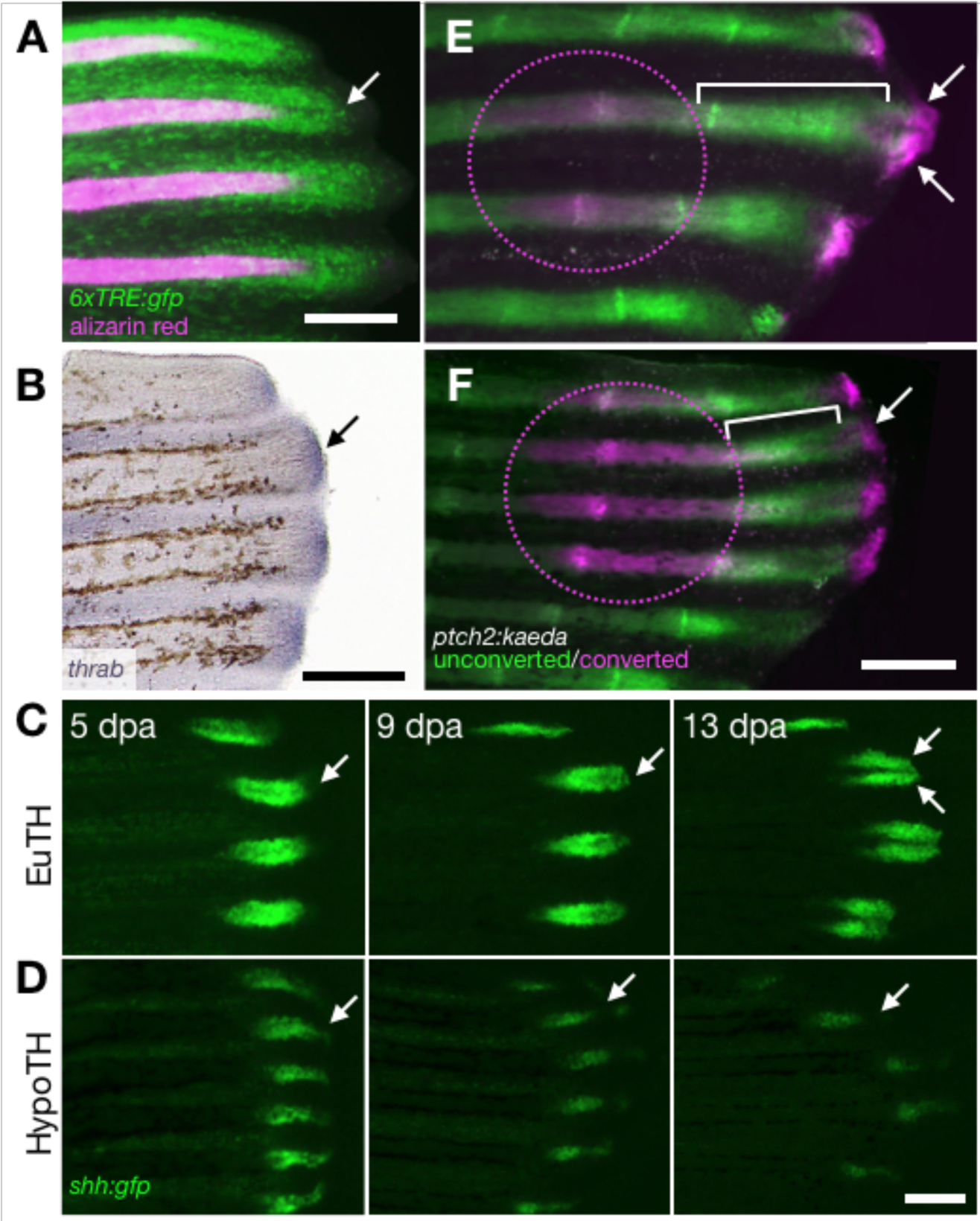
TH acts upstream of *shh* signaling to coordinate branching machinery with outgrowth. (**A**) Expression of TH activity reporter *6xTRE:gfp* in an 8 dpa euthyroid fish during branch morphogenesis; alizarin red shows mineralized bone. (**B**) *thrab* expression at the edges of a regenerating fin from a euthyroid prior to branch formation. (**C-D**) *shh:gfp* expression in regenerating fins of (C) euthyroid and (D) hypothyroid fish. In the fin of the euthyroid fish, branch is forming at 9 dpa and has formed by 13 dpa. (**E-F**) Regenerating fins of (E) euthyroid and (F) hypothyroid fish at 6 dpa, 48 hours following Kaeda photoconversion. Magenta circle shows photo-converted region. Unconverted and new Kaeda is green; photo-converted Kaeda is magenta. White brackets indicate the region of Kaeda produced since photoconversion; white arrows show displaced distal domains of converted Kaeda. In all images, arrows indicate second dorsal ray. Bars, 200 uM.

Since Shh signaling is essential for bifurcation (Quint et al., 2002; Zhang et al., 2012; Armstrong et al., 2017), we asked if TH was required for the Shh pathway. We found that *shh* was expressed in fin regenerates from hypothyroid fish, although the expression was somewhat attenuated, particularly during distal regeneration (Figure 4C-D). Notably, *shh* domains failed to separate in the absence of TH (Figure 4D). This contrasts with experimental treatments that can functionally block or delay bifurcation, but nonetheless show robust separation of *shh* expression domains (Azevedo et al., 2012; Armstrong et al., 2017).

Shh-active basal epidermal cells appear to undergo collective migration that directs osteoblast progenitors into lateral branches (Armstrong et al., 2017; Braunstein et al., 2020); we asked if this migrtion was dependent on TH. Using a photoconvertible *ptch2* reporter to read out Hh/Smo activity (Huang et al., 2012), we showed that even under hypothyroid conditions, Hh/Smo is active proximal to the tips of growing rays (brackets, Figure 4E-F). Moreover, TH was not required for previously Shh/Smo responsive cells to be displaced to the distal tips of growing rays (arrows, Figure 4E-F). In all, the molecular and cellular machinery underlying branch formation remains intact in hypothyroid fish, yet branching processes are not initiated. We conclude that TH acts upstream to deploy branching morphogenesis at the correct proximodistal location during outgrowth.

We have demonstrated that nuclear TH signaling is necessary and sufficient to coordinate axis patterning with growth of the fins, playing a regulatory or a permissive role to specify distal identity. Modulating TH titer fundamentally changes polarity of the appendicular skeleton independent of size, making the hormone an attractive target for fin diversification and changes to axis identity in other contexts.

## Materials and Methods

### Fish rearing conditions

Zebrafish were reared at 28°C with a 14:10 light:dark cycle, and fed 2-3 times per day. Fish stocks that were not TH-modulated (e.g. the *6xTRE:gfp* line and wild-type fish used for *in situs*) were fed with marine rotifers, *Artemia*, Gemma Micro (Skretting, Stavanger, NOR) and Adult Zebrafish Diet (Zeigler, Gardners PA, USA). To minimize potential introduction of exogenous TH, hypothyroid fish and euthyroid control fish were fed pure Spirulina flakes (Pentair, London, UK) instead of the enriched Zeigler diet.

### Fish lines

Wild-type zebrafish were of the *Tübingen* background. The following zebrafish mutants were used: hyperthyroid zebrafish were *opallus*^*b1071*^ (McMenamin et al., 2014); *longfin lof*^*dt2*^ (Van Eeden et al., 1996; Stewart et al., 2019); *shortfin sof*^*dj7e2*^ (Perathoner et al., 2014); *duox*^*sa9892*^ (Chopra et al., 2019); *thrab*^*vp31rc1*^ (Saunders et al., 2019). The following transgenic lines were used: *D. albolineatus Tg(tg:nVenus-v2a-nfnB)* and *D. rerio Tg(tg:nVenus-v2a-nfnB)* (McMenamin et al., 2014); *Tg(6xTRE-bglob1:eGFP)* (Matsuda et al., 2017); *Tg(−2*.*7shha:GFP)* (Neumann and Nuesslein-Volhard, 2000); *TgBAC(ptch2:Kaede)* (Huang et al., 2012).

### Thyroid follicle ablations

To ablate the thyroid follicles of *tg:nVenus-2a-nfnB D. rerio* or *D. albolineatus*, we incubated 4-5 dpf larvae overnight in 10 mM metronidazole (Mtz) with 1% DMSO, or with 1% DMSO alone as control in 10% Hanks, as in (McMenamin et al., 2014). Larvae were screened after ablation to ensure that no *nVenus* expression remained. Follicles do not regenerate after ablation, and thyroid-ablated zebrafish are functionally hypothyroid throughout their lives (McMenamin et al., 2014).

### Imaging

Zebrafish were anesthetized with tricaine (MS-222, ∼0.02% w/v in system water) and caudal fins were imaged on an Olympus SZX16 stereoscope using an Olympus DP74 camera, or on an Olympus IX83 inverted microscope using a Hamamatsu ORCA Flash 4.0 camera. Identical exposure times and microscope settings were used to compare experimental treatments, as well as capture fluorescent image series of the same fin over multiple days. Images were processed in cellSense (Olympus, Tokyo, JPN) and adjusted for contrast, brightness and color balance using GIMP and Keynote (Apple, Cupertino CA, USA); corresponding adjustments were made for images of both control and experimental fish. For paired images shown in figures, adult fish were size-matched to within 1 mm standard length (SL), measured as in (Parichy et al., 2009).

### Fin ray morphology quantifications

A landmark-based approach was used to quantify fin ray morphology from images using the R package StereoMorph (Olsen and Westneat, 2015). Landmarks were manually placed on fin ray segment joints and branch points for dorsal rays 1, 2, 3 and 4 and for ventral rays 1, 2, 3 and 4. The dorsal and ventral-most rays do not form bifurcations, and were omitted from subsequent analyses. Fin ray morphology was calculated using the pixel coordinates of these landmarks, and converted to millimeters.

While we demonstrated extremely strong effects of hypothyroidism on the position of the branches (see Figure 1 and Figure 1—Figure Supplement 1), we note that our analyses focused exclusively on the most lateral rays (D2-D4 and V2-V4), which were the most “distalized” of the rays in hypothyroid fish. The effects were even more profound on the medial rays in the fin, which we have never observed to form branches in hypothyroid backgrounds; therefore our focus on the lateral rays actually greatly decreases the effect size of our results and the morphological effects we demonstrate are considered conservative.

The placement of the bifurcation along the lengths of rays was analyzed as a function of number segments to the bifurcation, distance to the bifurcation and the proportion of the ray to bifurcation (i.e. [total distance from the body to the bifurcation]/[total ray length]). Rays grow continuously throughout the lifetime of the fish (see Figure 1—Figure Supplement 1H), and new segments are added distally. Thus, while the number of segments to bifurcation and the distance to bifurcation remain constant over adult growth, the proportion of the ray at bifurcation decreases with adult growth (Figure 1—Figure Supplement 1D). Therefore, when assessing the proportion of ray at bifurcation, we only considered fish 18-23 SL, excluding the largest and smallest individuals in the dataset. Rays that did not bifurcate were counted as having a proportion of ray at bifurcation of 1.0.

Note that non-branching rays are excluded from analyses of distance at bifurcation and number of segments at bifurcation. As many rays in hypothyroid backgrounds do not branch, particularly in smaller fish or in regenerated fins, these sample sizes for hypothyroid are frequently low and sometimes do not differ significantly from the distances or segment numbers of euthyroid controls (e.g. Figure 3—Figure Supplement 2 E-F). This reflects an altogether failure of the rays to branch, and the difference between bifurcating and non-bifurcating is better reflected in the comparisons showing proportion of ray at bifurcation, where non-branching rays are measured as 1.0.

### Statistical analysis

All analyses were performed in R v. 3.4.1 (R Core Team, 2016). Morphological data were analyzed by ANOVA, with ray identity and individual nested within treatment groups; significant differences between groups were then assessed with Tukey’s Honest Significant Differences test. In graphs showing few comparisons, significance is indicated as follows: *p* < 0.01, *; *p* < 0.001, **; *p* < 0.0001, ***. Marginal *p* values larger than 0.01 are listed in figures. In graphs showing many comparisons, significance is indicated on graphs by letter groups, with statistically indistinguishable groups sharing the same letter (based on a significance cutoff of 0.01, to account for multiple comparisons). Details of all statistical results available upon request.

Since proportion of ray length at primary bifurcation changes as fish grow (see Figure 1—Figure Supplement 1D), analysis of proportion at bifurcation are limited to fish 18-23 SL. Likewise, width of segments increases as fish grow, so analysis of segment width ratios are limited to fish 18-23 SL.

### Micro-computed tomography scans

Samples were fixed with 4% paraformaldehyde and embedded in 1% agar. Scanning was performed on a SkyScan 1275 high resolution micro-CT system (Bruker, Billerica MA, USA) at a scanning resolution of 10.5 um with an x-ray source voltage and current of 40 kV and 250 mA respectively. >2800 projection images were generated over 360° with a 0.1° rotation step and 6 averaging frames. Thresholding, ring artifact reduction, and beam hardening corrections were consistent across all scans during reconstruction using NRecon (Bruker). Reconstructed BMP slices were analyzed using Amira 6.5 (Thermo Fisher Scientific, Waltham MA, USA). Density heatmaps were generated with the volume rendering module and physics load transfer function. Density along the proximodistal axis of the 2^nd^ dorsal ray was measured from greyscale density renderings using ImageJ v. 1.49 (Schneider et al., 2012).

### RNAseq

Intact caudal fin tissue was sampled from sibling adults (>18 SL) reared under euthyroid or hypothyroid conditions. Each fish was first anesthetized with tricaine (MS-222, ∼0.02% w/v in system water) and the entire caudal fin was amputated using a razor blade. Proximal, middle and distal regions of the fin were sampled immediately after and flash frozen in a dry ice-ethanol bath. Three biological replicates each containing five fin regions were collected for both TH backgrounds. RNA was extracted the same day with Zymo Quick-RNA Microprep kit R1050 (Zymo Research, Irvine CA, USA). Sample libraries were made with NEBNext Ultra II RNA Library Prep kit and sequenced on an Illumina Novaseq 6000 platform with 20M 150bp paired-end sequences per sample. Quality check, library preparation and sequencing were done by Novogene. Raw sequence reads were aligned to Zebrafish GRCz11 using STAR version 2.7.3 (Robinson et al., 2010), gene counts were called using Ensembl GRCz11 gene annotation. Differential gene expression analyses were performed with Bioconductor package edgeR (Robinson et al., 2010). Genes with a log_2_ fold change higher than 2 and a false discovery rate lower than 0.01 were considered significant. Gene Ontology analysis was performed using clusterProfiler (Yu et al., 2012).

### Computational model

To explore the phenotypes that could be theoretically produced by changing proximodistal identity relative to size, we produced a simple computational model in R (R Core Team, 2016). The “size factor” input determined the length of the starting segment, from which all other segment lengths were calculated. The “distalization factor” input determined both the location of the branch and the rate at which segments shortened. Our goal with this model was not to perfectly recapitulate fins observed; rather to demonstrate the diversity that may theoretically be generated by decoupling and independently modulating theoretical processes regulating size and distalization.

### Fin amputation

All whole-fin regeneration experiments were performed on adult zebrafish 18-23 mm SL. Caudal fins were amputated from anesthetized fish under a stereoscope at the 4-5^th^ ray segment using a razor blade. Lateral ray excisions (Figure 4—Figure Supplement 1B-E) were performed with a razor blade, and continuously trimmed every second day at the base of the ray to prevent lateral ray regeneration.

### Drug treatments

Working stocks of L-thyroxine (T4; Sigma-Aldrich, St. Louis MO, USA) were diluted in 0.5 M NaOH, then diluted into fish room water to a final concentration of 10 μM T4 with 194 nM NaOH. Vehicle controls were 194 nM NaOH in fish room water. For treatments longer than 24 hours, 90% water changes were performed 6 days a week throughout the treatment period.

### Quantitative PCR

Regenerating fin tissue (∼4 segments wide) was sampled on days 5, 7, 9 and 15 post amputation in adult EuTH zebrafish. Four samples were pooled as one biological replicate and a total of three biological replicates were collected at each time point. Sample was stored in RNAlater at -80°C. RNA was extracted with Quick-RNA Microprep kit (Zymo Research, Irvine, CA) cDNA was produced using the SuperScript IV First-Strand Synthesis System (Invitrogen, Carlsbad, CA), qPCR of *thrab* was performed on QuantStudio 3 (Applied Biosystems, Foster City CA, USA) cycler using primers F:tctgatgccatcttcgacttg; R: gtacatctcctggcacttctc. Data was analyzed with ThermoFisher Connect software (Thermo Fisher Scientific).

## Acknowledgements

For valuable input, discussion and assistance, we thank Andy Aman, Kunal Chopra, Matt Gregas, Andrea Kirmaier, Kate McCusker, Andrew Wagner and Andrew Williston and all members of the McMenamin Lab. For generously sharing lines, we thank Enrique Amaya, James Cooper, Matthew Harris, Rolf Karlstrom, Jennifer Lanni, David Parichy, Dedier Stanier, Kryn Stankunas and their respective labs.

---

| Funder | Grant reference number | Author |
| --- | --- | --- |
| National Science Foundation | CAREER 1845513 | Sarah McMenamin |
| National Institutes of Health | R00GM105874 | Sarah McMenamin |
| National Institutes of Health | R03HD091634 | Sarah McMenamin, Patricia Hernandez |
| Charles A. King Trust<br>Postdoctoral Fellowship |  | Yinan Hu |
| National Institutes of Health | R35GM122471 | David Parichy |

## Competing interests

None.

## Data and material availability

Requests for materials and complete datasets should be addressed to Sarah McMenamin.

## Figure Supplements

**Figure 1−Figure Supplement 1.**
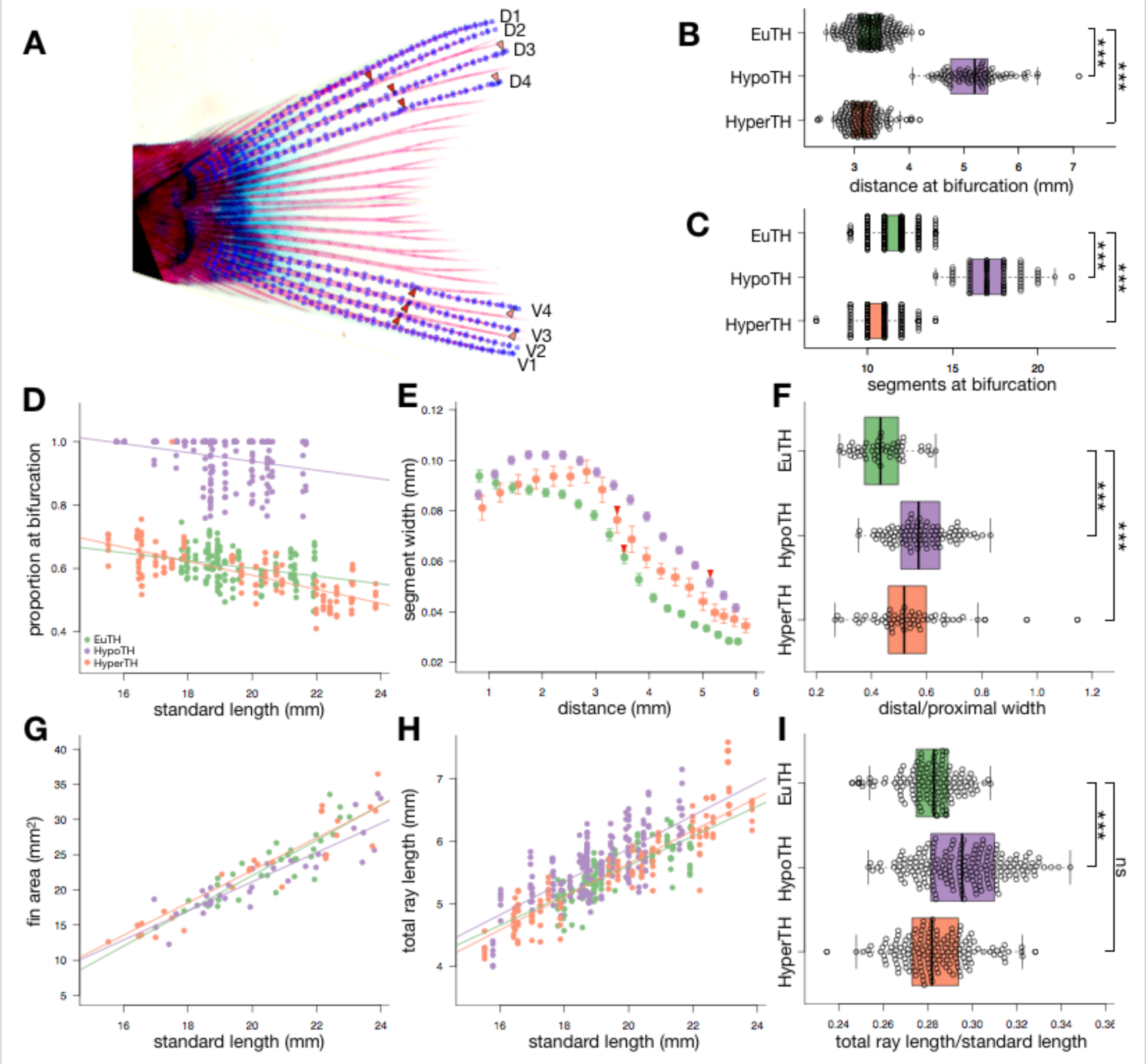
TH regulates proximodistal morphology independent of fin area or ray length. (**A**) Example of landmarks placed on a single fin during digitization. For dorsal rays 1-4 (D1-D4) and ventral rays 1-4 (V1-V4), landmarks were placed at the joints between segments to measure segment length; two landmarks were placed across the middle of each segment to measure width. Landmarks were placed at each bifurcation node (indicated with red arrowheads). The distance to the bifurcation on each ray was the sum of all segments (and the partial segment) proximal to the primary bifurcation. The total length of each ray was the sum of all the complete segments in the ray following the dorsalmost (for dorsal rays) or ventralmost (for ventral rays) bifurcation of the ray. The distalmost segment was not counted in the total length. Box plots showing (**B**) distance to the caudal peduncle from the bifurcation, (**C**) number of segments proximal to the bifurcation. (**D**) Plot showing proportion of the total fin ray length at the bifurcation as a function of standard length of the fish. (**E**) Plot showing the average width of each segment (starting at the 3rd segment from the body) as a function of the distance from the segment to the caudal peduncle. Error bars represent 95% confidence interval. Red arrowheads indicate the average location of the primary bifurcation in each background. (**F**) The ratio of distal segment width (segments 15-17) to proximal segment width (segments 5-7). (**G**) Plot showing area of the fin as a function of standard length of the fish. (**H**) Plot showing length of each ray as a function of standard length of the fish. (**I**) Total ray lengths standardized by standard length of the fish. For all plots, green, euthyroid; purple, hypothyroid; orange, hyperthyroid. Significance determined by ANOVA followed by Tukey’s HSD.

**Figure 1−Figure Supplement 2.**
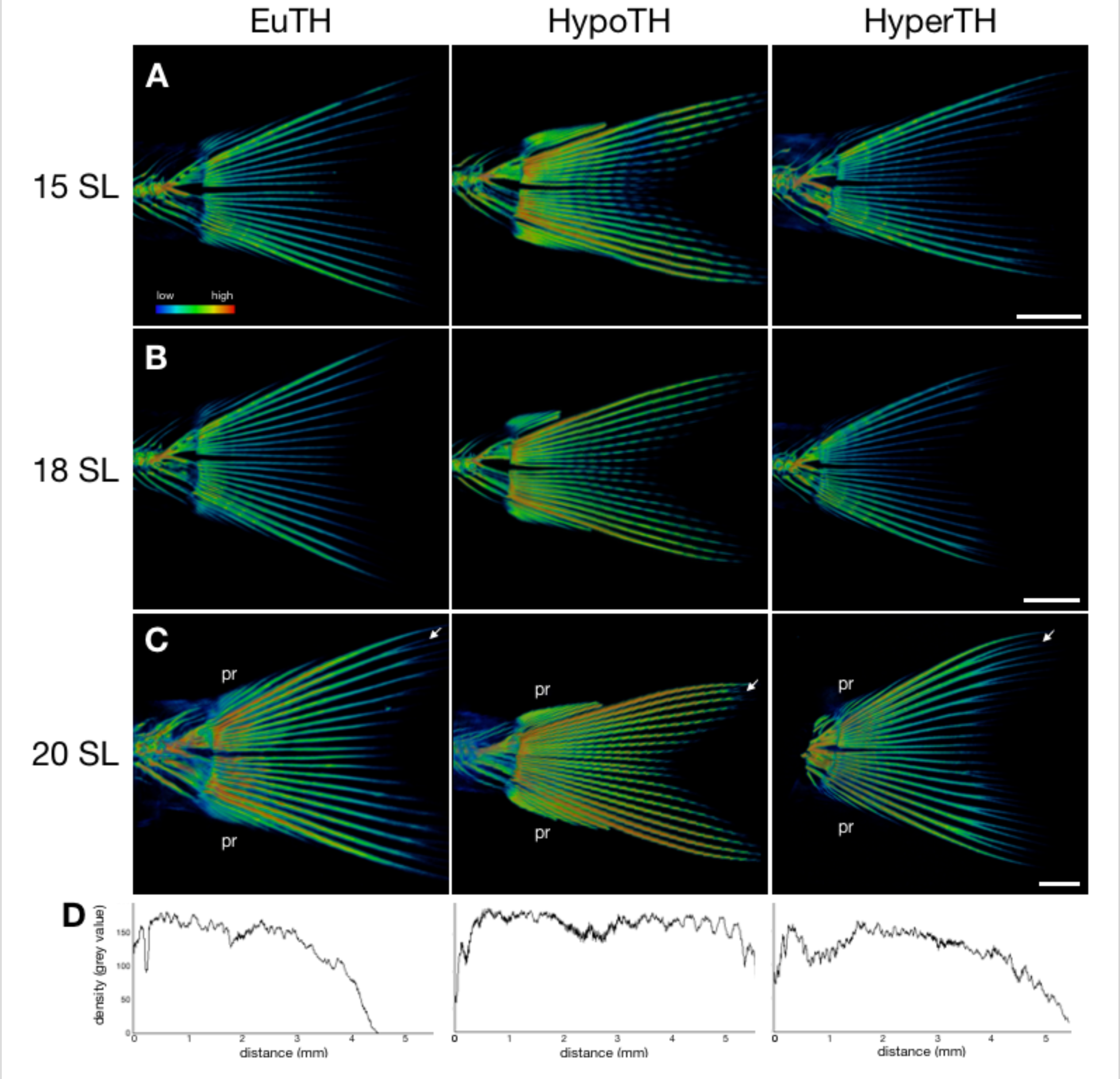
TH regulates density of fin rays along the proximodistal axis. (**A-C**) Micro-CT scans of fins from fish reared under different TH profiles, at three different standard lengths. Scanning conditions and thresholds are identical within SL groups. Warmer colors show higher density tissue; cooler colors show less dense tissue. White arrows indicate rays used to generate density graphs in (D). Scale bars, 1 mm. pr, procurrent rays. Note that procurrent ray density appears to scale inversely with TH. (**D**), Density of the second dorsal ray of 20 SL individuals along the proximodistal axis. Note that ray from the hypothyroid background retains high density along the length of the ray and into distal regions.

**Figure 1−Figure Supplement 3.**
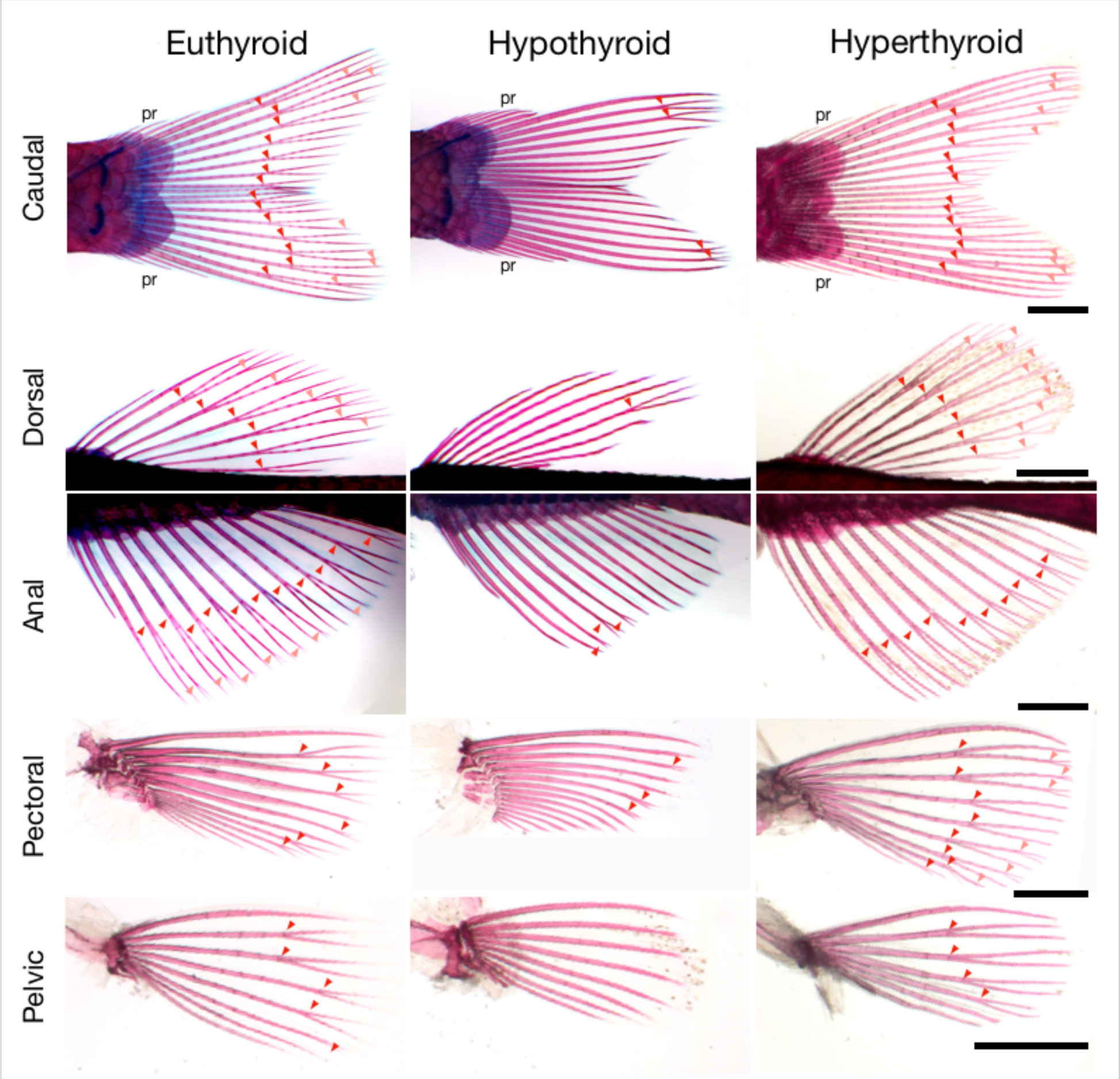
TH is required for distal ray morphology in paired and medial fins. The three medial fins (caudal, dorsal and anal) and two paired fins (pectoral and pelvic) of adult zebrafish reared under three TH profiles, cleared and stained. Red arrowheads indicate primary bifurcations; pink arrowheads indicate secondary bifurcations. pr, procurrent rays. Note that the number of procurrent rays scales inversely with developmental TH titer: procurrent rays in euthyroid = 5.2; standard deviation = 0.4. Procurrent rays in hypothyroid fish are more longer, more robust and more numerous (ave. = 7.2; SD = 0.7); hyperthyroid fish have fewer, shorter procurrent rays (ave. = 4.6; SD = 0.5). Scale bars, 1 mm.

**Figure 1−Figure Supplement 4.**
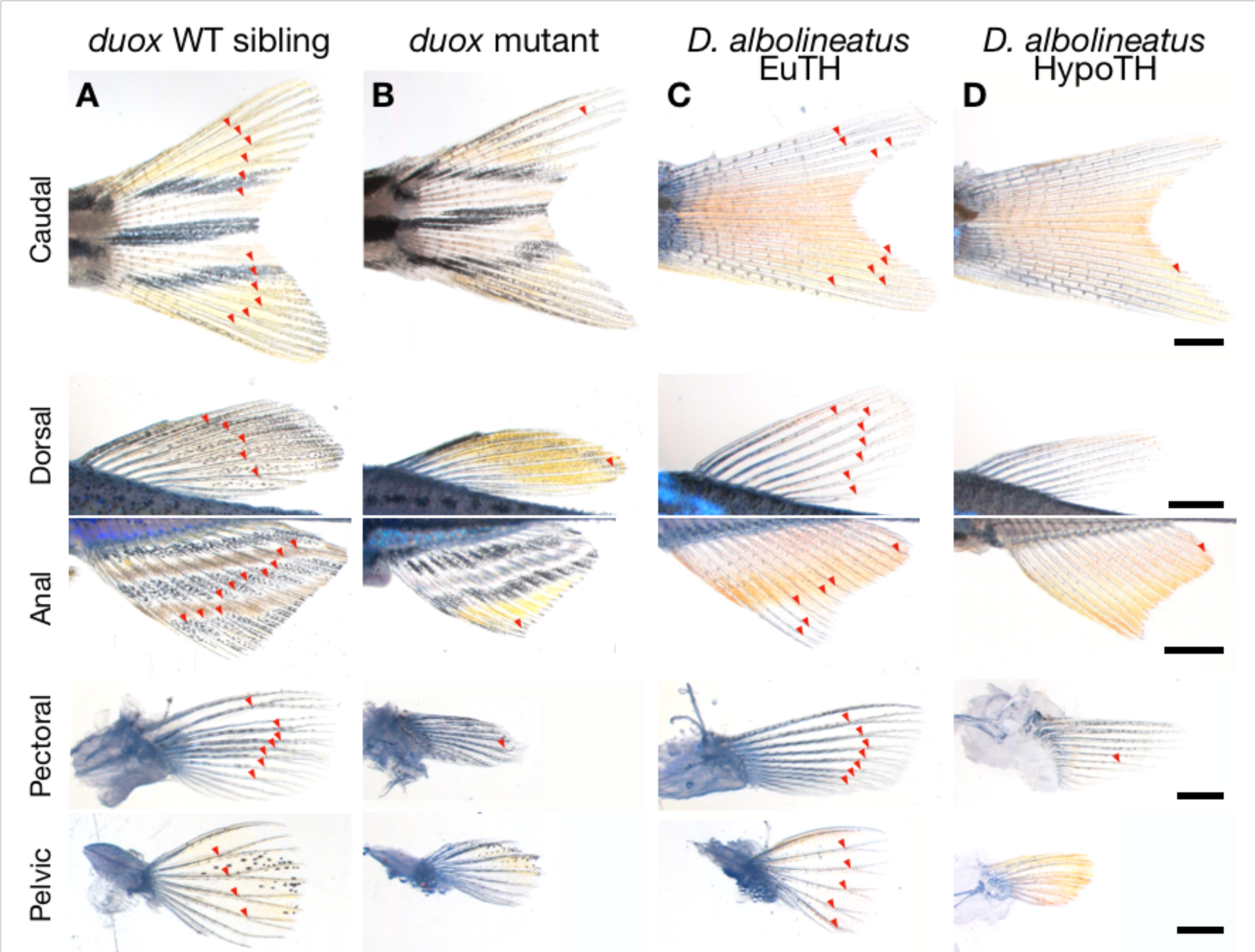
Hypothyroid zebrafish mutant and hypothyroid *Danio albolineatus* show proximalized fins. (**A-B**) Fins of wild-type sibling and *doux* mutant zebrafish. (**C-D**) *Danio albolineatus* siblings reared under either euthyroid or hypothyroid conditions. Arrowheads indicate bifurcations. Scale bars, 1 mm.

**Figure 1−Figure Supplement 5.**
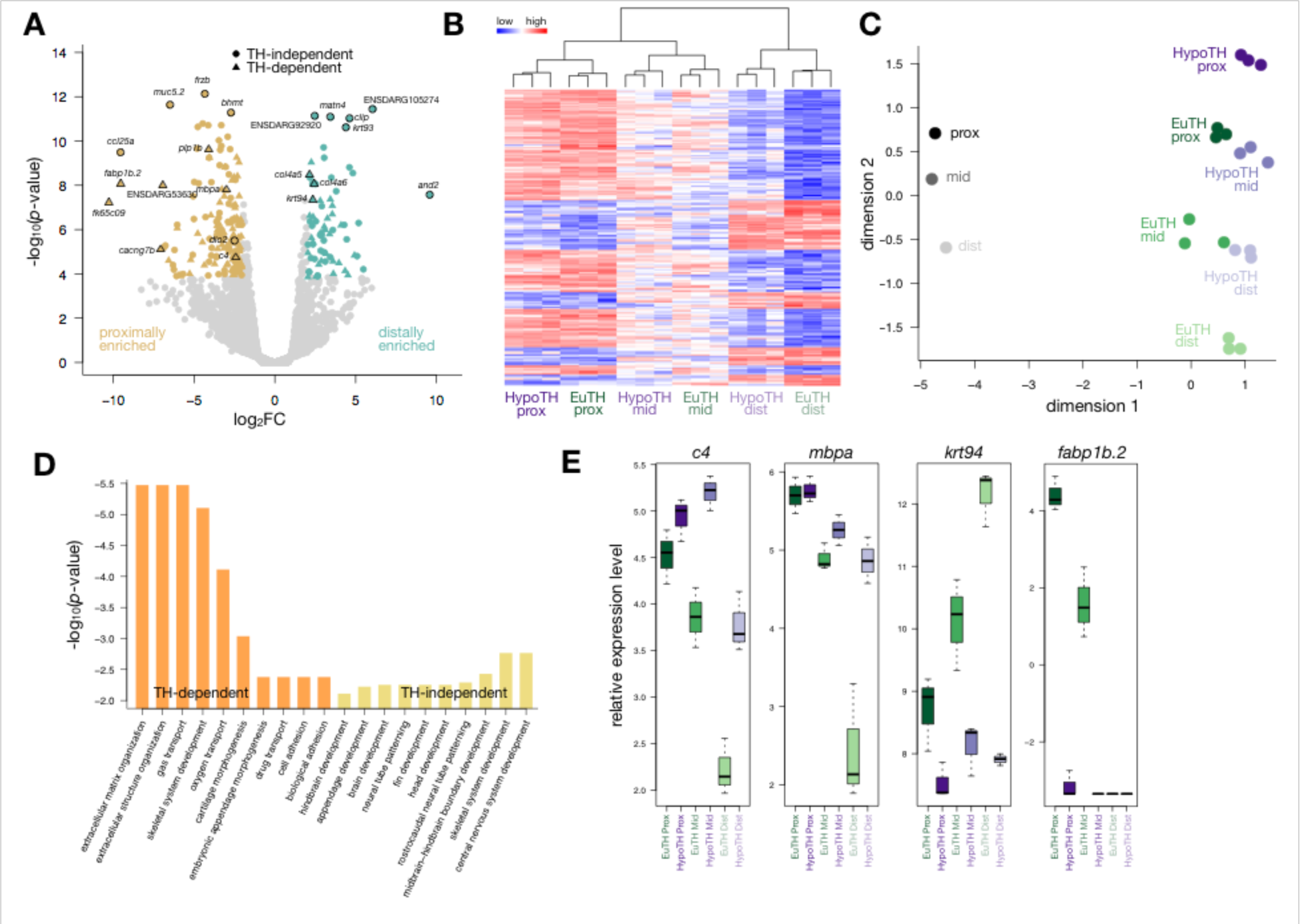
TH promotes distal gene expression profiles. (**A**) Volcano plot showing genes differentially expressed between proximal (yellow) and distal (blue) regions of fins from euthyroid zebrafish. Genes with a proximodistal gradient independent of TH are shown as circles; genes with a proximodistal gradient dependent on TH are shown as triangles. Note that *dio2*, which encodes an enzyme that converts TH into its more active form, is proximally enriched in a non-TH-dependent manner. *fabp1b*.*2* is associated with RA metabolism (Rabinowitz et al., 2017), and is proximally enriched, also showing TH-dependence. (**B**) Heat map of the 233 genes expressed in a distal gradient in euthyroid fins; the subset of these genes that are show TH-dependence for gradient expression are shown in Fig. 1G. Dendrogram at top represents hierarchical clustering of the samples. Relative high expression shown as red; relative low expression shown as blue. (**C**) Multidimensional scaling plot comparing gene expression profiles of different regions of the fin from eu- and hypothyroid fish; each data point represents a single biological replicate. Also shown is a previously published reference dataset (greyscale; Rabinowitz et al., 2017), with which our dataset shows high consistency. Note that dimension 2 captures proximodistality of all the datasets, and that hypothyroid transcriptomes are overall shifted ‘proximally’ along this axis relative to their euthyroid controls. (**D**) Gene Ontology terms for genes that show proximodistal expression gradients. Terms enriched in gradients that are dependent on TH shown in orange; terms enriched in gradients retained in hypothyroid contexts are shown in yellow. (**E**) Boxplot comparing expression levels of selected proximodistally-differential genes in different fin regions between eu- and hypothyroid fish. Each of these four genes was identified in Rabinowitz (2017) as proximally or distally enriched by both RNAseq and proteomic analysis. Note that while *c4, mbpa* and *krt94* show proximalized expression patterns in fins of hypothyroid fish, *fabp1b*.*2* expression is merely depressed at all proximodistal levels.

**Figure 2−Figure Supplement 1.**
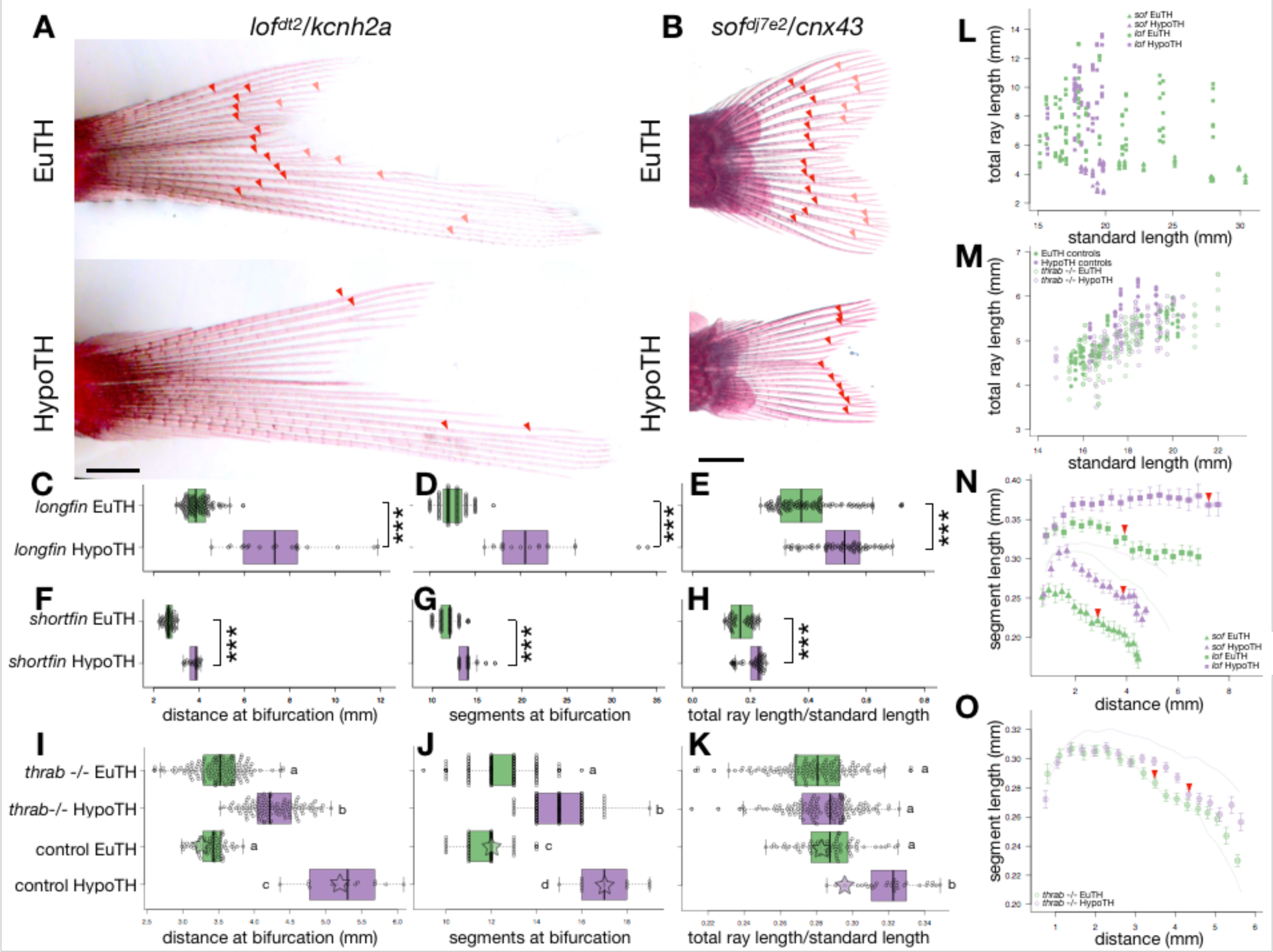
TH distalizes fins in *shortfin* and *longfin* mutant backgrounds, *thrab* positively and negatively regulates proximodistal identity. **(A-B)** Caudal fins from *longfin* and *shortfin* mutants reared under euthyroid or hypothyroid profiles, cleared and stained. Red and pink arrowheads indicate primary and secondary bifurcations, respectively. Scale bars, 1 mm (**C-K**) Boxplots showing (C, F, I) distance from the body to the primary bifurcation; (D, G, J) the number of segments proximal to the bifurcation; and (E, H, K) the total length of the ray standardized by the standard length of the body. In I-K, data are shown for control non-mutant eu- and hypothyroid reared side-by-side and under identical conditions to the *thrab* -/-mutants. For comparison, overlaid stars indicate the mean values for all non-mutant eu- and hypothyroid (i.e. those shown in Figure 1 and Figure 1—Figure Supplement 1). Significance determined by ANOVA followed by Tukey’s HSD. (**L-M**) Length of rays as a function of standard length. (**N-O**) Plots showing the average length of each segment (starting at the 3rd segment from the body) as a function of the distance from the body to that segment. Red arrowheads indicate the average location of the primary bifurcation in each background. For reference, the average lengths for non-mutant eu- and hypothyroid (i.e. data plotted in Figure 1F) are shown as light lines.

**Figure 2−Figure Supplement 2.**
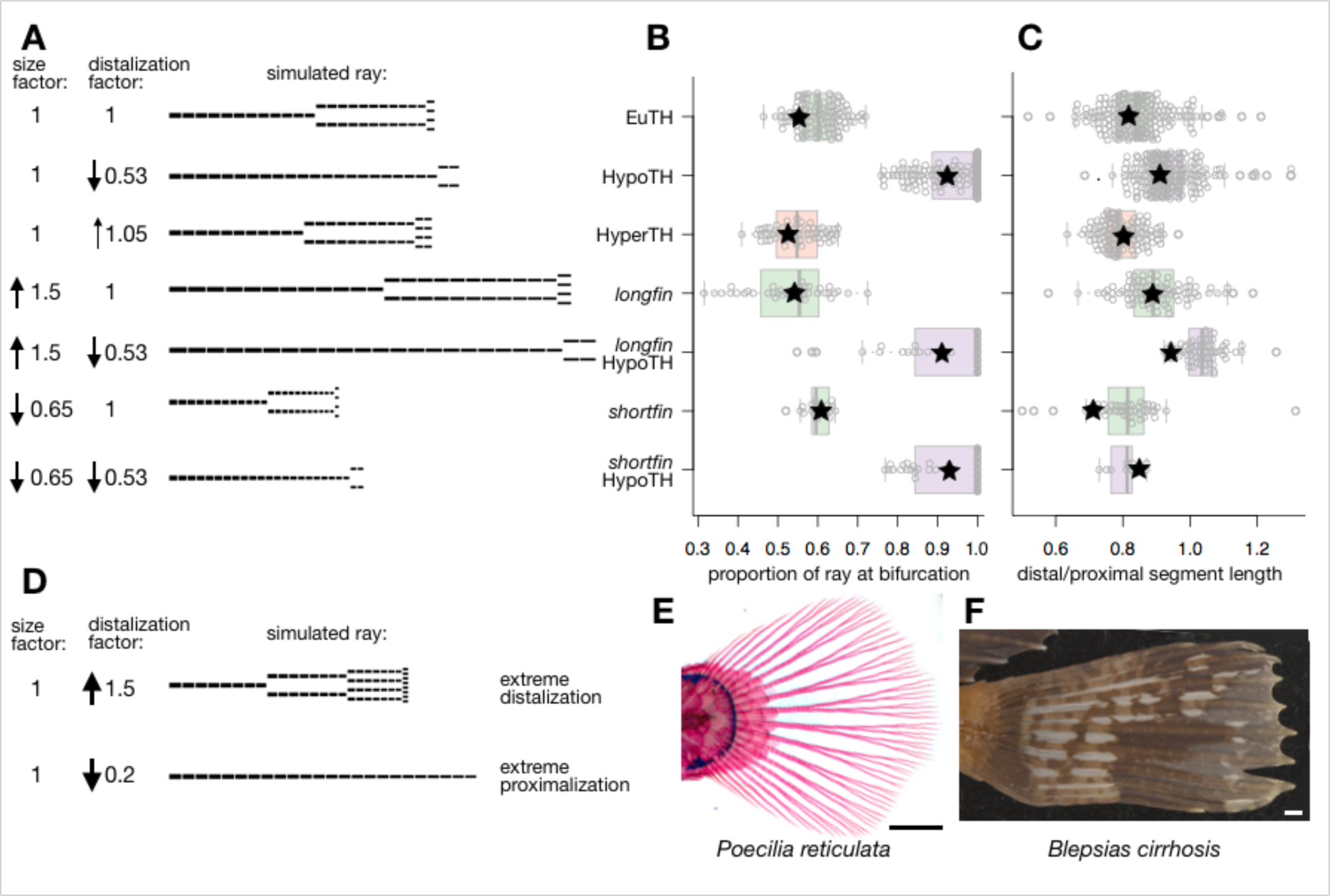
A computational model generates fin ray diversity by independently modulating size and distalization. (**A**) Computational model simulating segmented fin rays. Up and down arrows represent the direction and magnitude of relative changes, which are also given numerically. Note that the model accurately predicts that proximalized rays will be slightly longer due to the lengthened distal segments. (**B-C**) Proportions of virtual rays (black stars) corresponding to the inputs from A. For comparison, these values are overlaid on the proportions from actual rays in different backgrounds. (**D**) Extreme relative increase or decrease in the distalization factor simulated rays that coarsely resemble rays of (**E**) guppy *Poecilina reticulate* or (**F**) silver spotted sculpin *Blepsias cirrhosus*. Scale bars, 1 mm.

**Figure 3−Figure Supplement 1.**
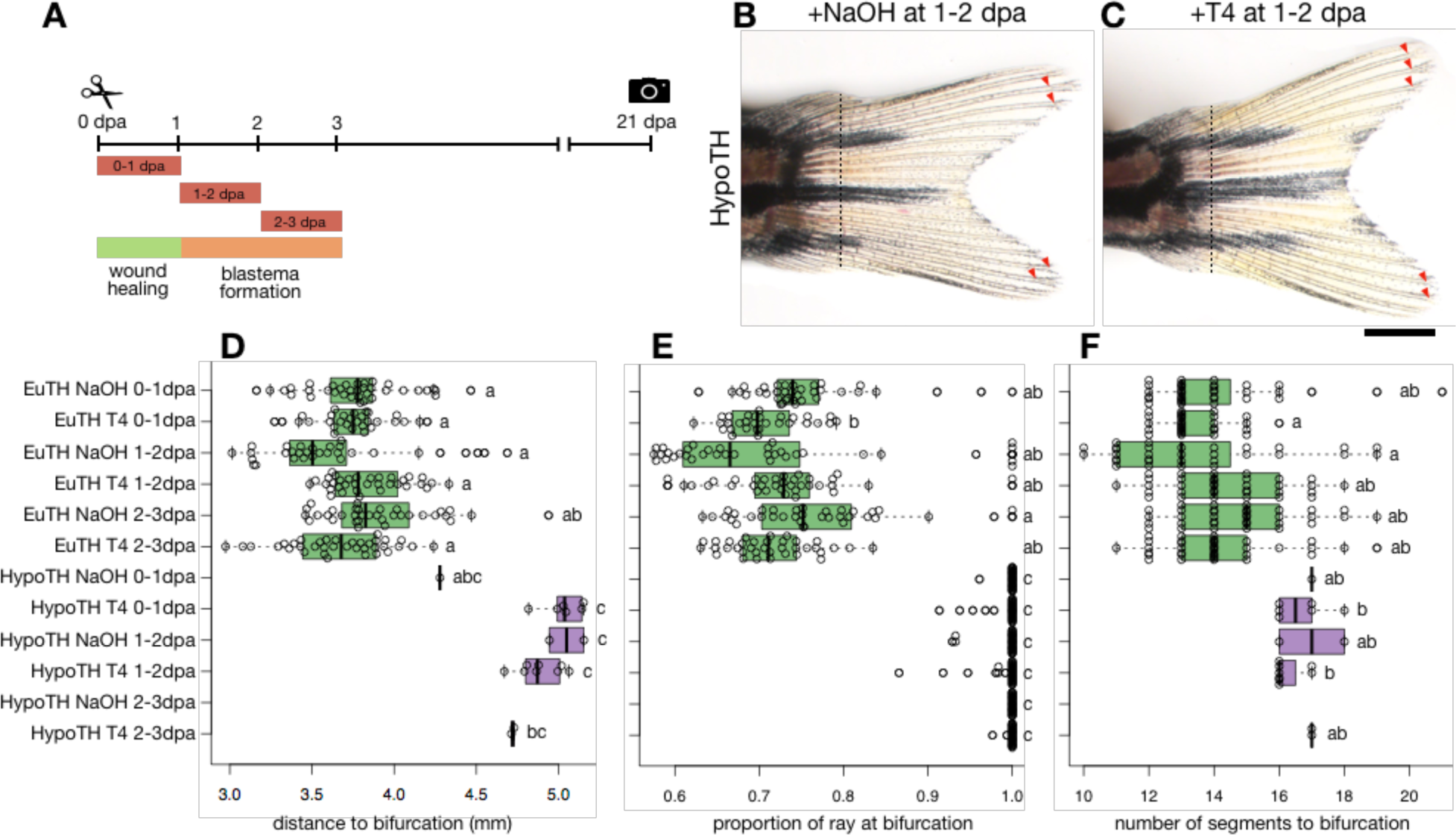
Treating the blastema with exogenous TH does not rescue distalization in regenerate. (**A**) Experimental treatment timeline. Fish were treated with either 10uM T4 or NaOH vehicle for periods 0-1, 1-2 or 2-3 dpa. (**B-C**) Fins from hypothyroid fish treated with either NaOH vehicle or 10uM T4 for 24 hours starting 1 day after amputation (1-2 dpa treatment). Dashed lines indicate planes of amputation. Arrowheads, bifurcations; Bar, 1 mm. (**D**) Distance from body at bifurcation, (**E**) Proportion of total ray length at bifurcation, (**F**) Number of segments proximal to bifurcation. Note that non-bifurcating rays are not represented in D and F, and therefore the sample size is extremely small (or nonexistent) for NaOH-treated hypothyroid fish because very few of these rays form branches. Significance groups determined by ANOVA and Tukey’s HSD.

**Figure 3−Figure Supplement 2.**
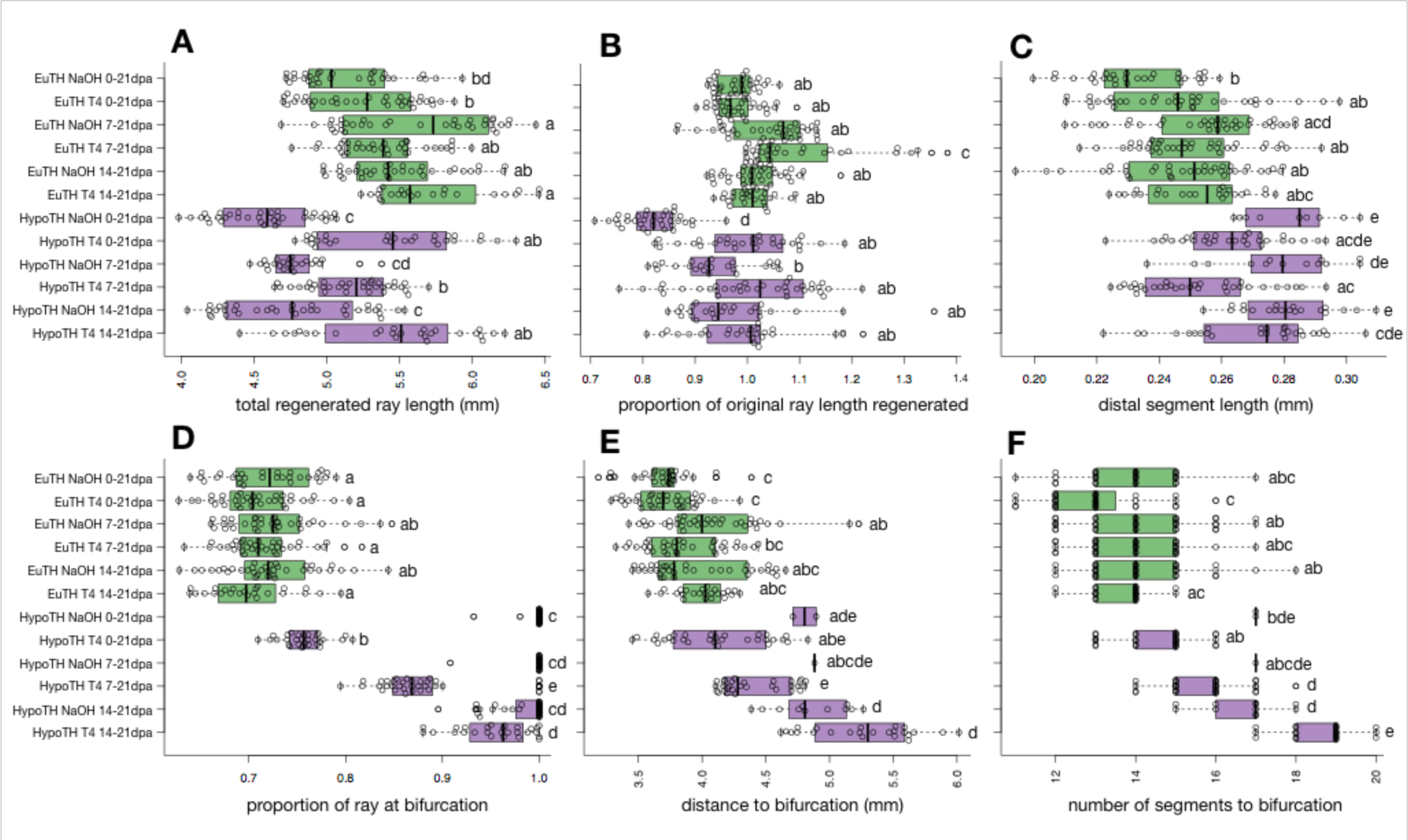
TH treatment during regeneration rescues distal morphologies in fins of hypothyroid fish. (**A**) Total length of regenerated rays. (**B**) Proportion of original length of ray regenerated (total length before amputation/total length after regeneration). Note that fins of hypothyroid fish regenerate more slowly than euthyroid counterparts, and that a euthyroid regeneration speed is rescued with T4 treatment. (**C**) Average length of distal segments 15-17. (**D**) Proportion of total ray length at bifurcation. (**E**) Distance from body at bifurcation. (**F**) Number of segments proximal to bifurcation. Note that non-bifurcating rays are not represented in E and F, and therefore the sample size is extremely small for NaOH-treated hypothyroid fish because very few of these rays form branches. Significance groups determined by ANOVA and Tukey’s HSD.

**Figure 4−Figure Supplement 1.**
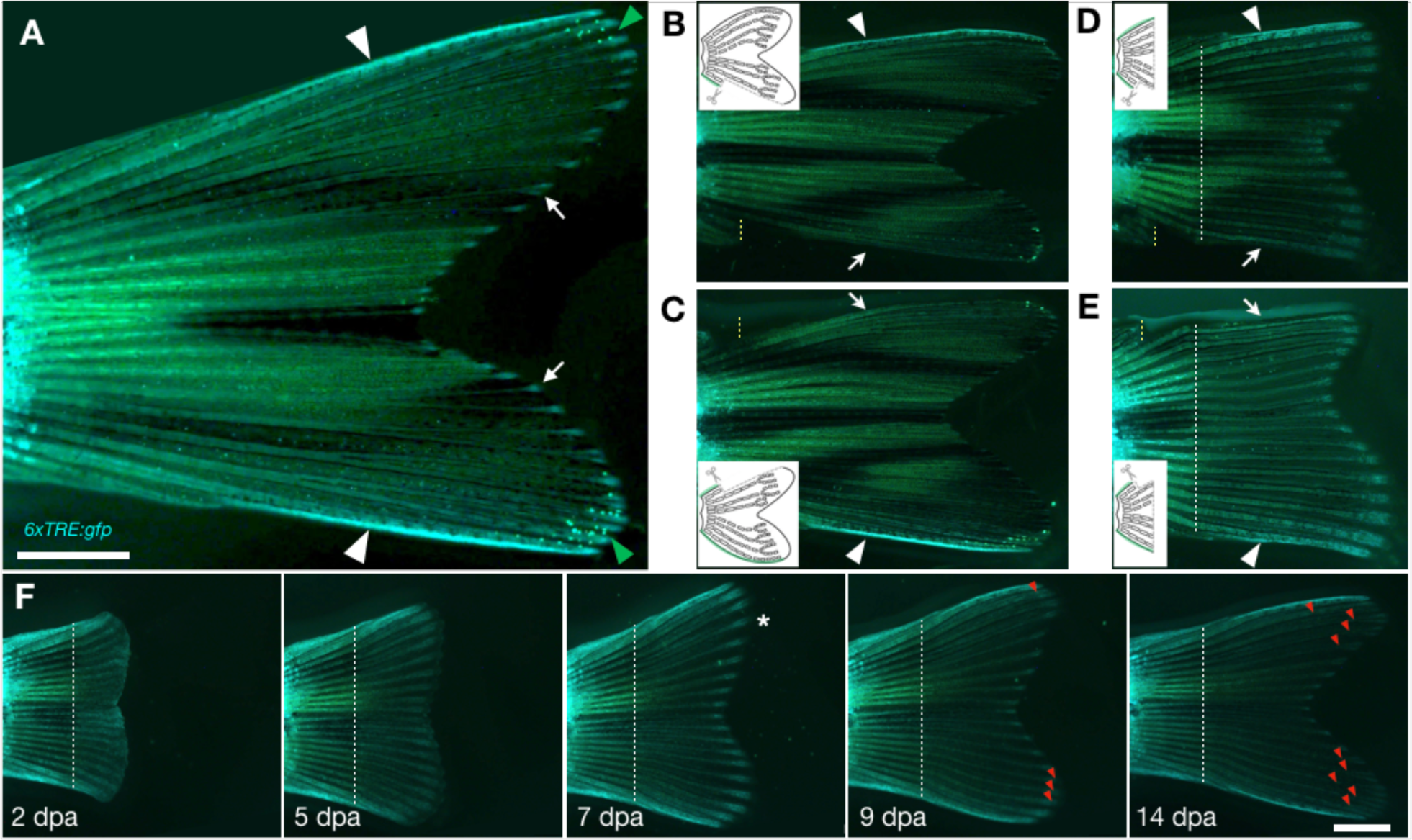
TH activity in lateral tissues and at the tips of uninjured and regenerating fin rays in euthyroid fish. (**A**) *6xTRE:gfp* expression in uninjured fin. White arrowheads, TH activity in tissue adjacent to lateral rays. Arrows, TH activity at the distal tips of rays. Green arrowheads, leucophores at the lateral tips of the rays. Note GFP appears blue-green, while autofluorescent tissues and pigment cells appear yellow-green. (**B-C**) *6xTRE:gfp* expression in fins at 8 days after amputating the (B) dorsal or (C) ventral rays. Note that the next most lateral rays, D2 and V2, do not adopt the high TH activity when their more lateral neighbor is removed. (**D-E**) Expression in fins at 8 days after a full amputation (white dashed line) followed by repeated trimming of the (D) dorsal or (E) ventral lateral rays (yellow dashed line). Again, note that D2 and V2 do not adopt high TH activity even when regenerating in the absence of a lateral neighbor. In B-E, arrowhead indicates uninjured or regenerated lateral ray; arrow indicates side from which the lateral ray was amputated. White dashed lines indicate plane of amputation; yellow dashed lines indicate plane along which lateral rays were trimmed. (**F**) A single individual showing the course of *6xTRE:gfp* expression during regeneration. Asterisk indicates a spatiotemporal pulse of TH activity preceding branch formation. Dashed lines indicate amputation planes; red arrowheads indicate bifurcation nodes as determined by matched brightfield images. Scale bars, 1 mm.

**Figure 4−Figure Supplement 2.**
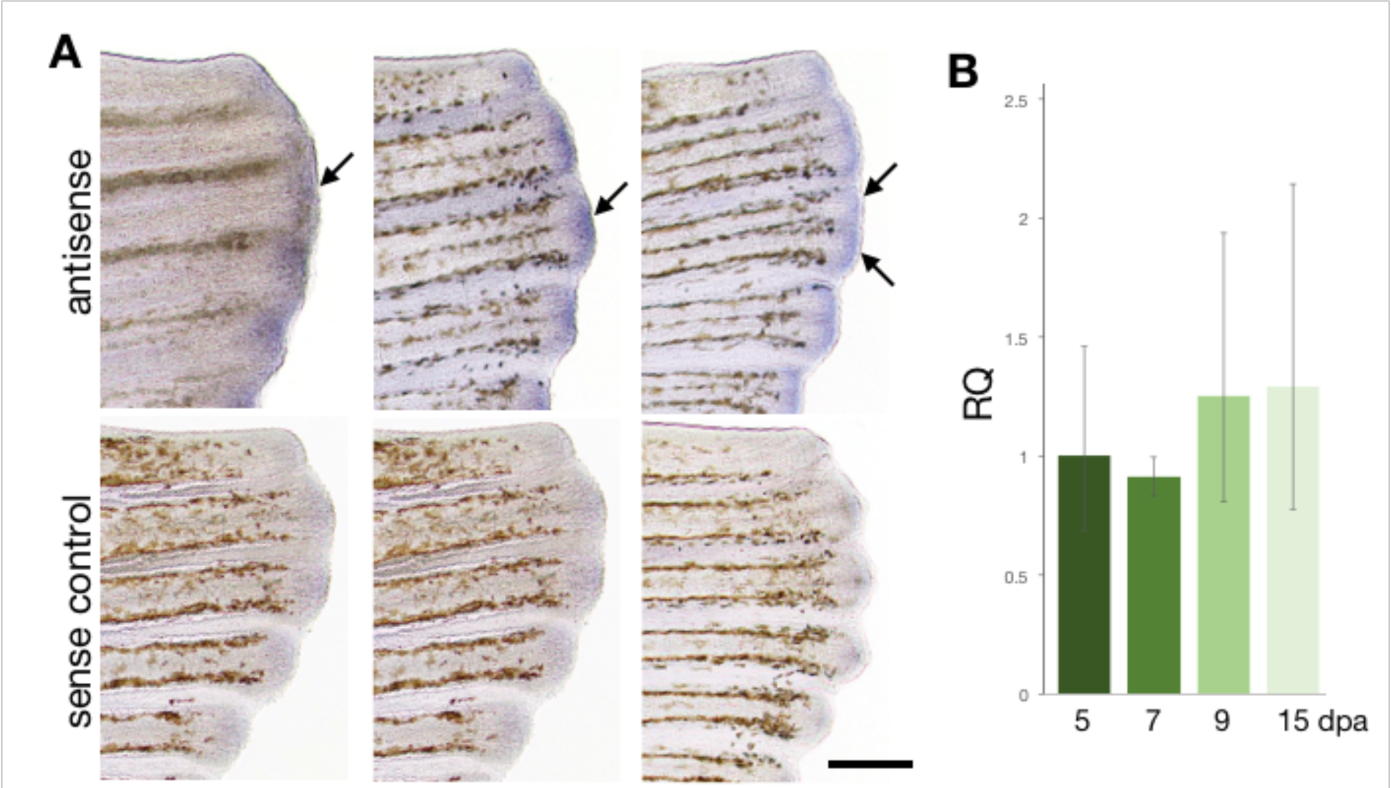
*thrab* is expressed at multiple stages of fin regenerative morphogenesis. (**A**) *In situ* hybridization of fins at different stages of bifurcation morphogenesis in wild-type fins. Arrows, *thrab* expression at the leading edges of growing rays. Top, antisense probe for *thrab*; bottom, sense control for *thrab*. (**B**) Relative quantification of gene expression levels from the distal edge of regenerating fins at 5, 7, 9 and 15 dpa.

## Notes

### Competing Interest Statement

The authors have declared no competing interest.

### Summary of Updates

Inclusion of Figure 3; updates to text.

